# Nucleus-Independent Transgenerational Small RNA Inheritance in *C. elegans*

**DOI:** 10.1101/2023.06.20.545749

**Authors:** Itai Rieger, Guy Weintraub, Itamar Lev, Kesem Goldstein, Dana Bar-Zvi, Sarit Anava, Hila Gingold, Shai Shaham, Oded Rechavi

## Abstract

Studies using *C. elegans* nematodes demonstrated that, against the dogma, animals can transmit epigenetic information transgenerationally. While it is now clear that in these worms ancestral RNA interference (RNAi) responses continue to regulate gene expression for many generations, it is still debated whether the primary agent that perpetuates heritable silencing is RNA or chromatin, and whether the information is communicated to the next generation inside or outside of the nucleus. Here we take advantage of the tractability of gene-specific double stranded RNA-induced heritable silencing to answer these questions. We demonstrate that RNAi can be inherited independently of any changes to the chromatin or any other nuclear factors via mothers that are genetically engineered to transmit only their ooplasm but not the oocytes’ nuclei to the next generation. Nucleus-independent RNA inheritance depends on ZNFX-1, an RNA-binding germ granule resident protein. We find that upon manipulation of normal germ granules functions (in *pptr-1* mutants) nucleus-independent RNA inheritance becomes stronger, and can occur even in *znfx-1* mutants. Utilizing RNA sequencing, chimeric worms, and sequence polymorphism between different *C. elegans* isolates, we identify endogenous small RNAs which, similarly to exogenous siRNAs, are inherited in a nucleus-independent manner. From an historical perspective, nucleus-independent inheritance of small RNAs might be regarded as partial vindication of discredited cytoplasmic inheritance theories from the 19^th^ century, such as Darwin’s “pangenesis” theory.

## Introduction

In the 19th century, before the chromosomal theory of inheritance, scientists argued over the role that the nucleus plays in transmission of information to the next generation. Hertwig, Strasburger, von Kolliker, and Weismann, hypothesized that the nucleus is the carrier of hereditary properties (Churchill, 1987; Weismann, 1893), while other contemporaries, most notably the Swiss botanist Carl Nägeli, believed that the heritable agents ignore cellular and sub-cellular boundaries (Nägeli claimed that the hereditary substance is somewhere in the “protoplasm”) (Rogers, 2014, pages 136-137). Similarly, Darwin believed in soma-to-germline inheritance of extra-nuclear information (via “gemmules’’, reviewed in: Bowler, 2003; Liu, 2008, 2018). In the second half of the 20th century, the understanding that DNA is the heritable material ended this discussion (although the DNA of the mitochondria and the chloroplast resides of course outside of the nucleus). Most studies on transgenerational epigenetic inheritance focused on nuclear inheritance of chromatin changes; Nevertheless, it remained unclear whether other types of epigenetic information (non-DNA encoded), such as small RNA-controlled responses, can be independently inherited in the cytoplasm.

Transgenerational epigenetic inheritance of environmental responses has been described in many organisms (reviewed in: Jablonka and Raz, 2009; Radford, 2018), but is especially well-understood in *Caenorhabditis elegans* nematodes; It is relatively straightforward to study transgenerational effects in *C. elegans*, as they exhibit gene-specific long term heritable RNA interference (RNAi) responses. Transgenerational RNAi entails the synthesis, processing and inheritance of exogenous short interfering RNAs (exo-siRNAs) following administration of exogenous double-stranded RNAs (dsRNAs) (Fire et al., 1998). In addition to being a useful tool for investigating gene functions, exogenous dsRNA-derived siRNA-mediated silencing is physiologically relevant, as it enables anti-viral defense, and as both animals and plants evolved mechanisms for taking up dsRNA from the environment and even from other organisms (Chen and Rechavi, 2022). In addition to exogenous dsRNA-induced siRNAs, worms respond to different environmental challenges by synthesizing endogenous small interfering RNAs (endo-siRNA) that regulate gene expression and affect their physiology across multiple generations (Anava et al., 2014; Ni et al., 2016; Rechavi et al., 2014, 2011). In comparison to experiments using gene specific dsRNA-induced silencing, transgenerational inheritance of endogenous small RNAs in response to stress is more challenging to study, as these complex responses reshape the entire transcriptome and entail many indirect interactions between numerous genes.

The dedicated machinery which enables RNAi inheritance in *C. elegans* has been extensively studied (Grishok et al., 2000; Alcazar et al., 2008; Gu et al., 2009; B. a Buckley et al., 2012; Grishok, 2013; Houri-Ze’evi et al., 2016; Wan et al., 2018; Houri-Zeevi et al., 2020), and heritable siRNAs were found to typically be 22 nucleotides long and to start with the nucleotide guanine (hereafter 22Gs). Inherited 22G small RNAs persist in the progeny despite the dilution of the parental RNAs, as they are synthesized anew in every generation by RNA-dependent RNA polymerases (RdRP) which use the target RNA as template (Sijen et al., 2001). RDE-3, a nucleotidyltransferase, adds untemplated polyUG sequences to the mRNAs of RNAi-targeted genes (Preston et al., 2019), marking them for additional rounds of amplification by RdRPs, and thus leading to the generation of more 22G to induce silencing and perpetuate inheritance (Shukla et al., 2020).

Are amplified small RNAs transmitted to the next generation inside or outside of the germline nucleus? Different studies examined the role of different cellular compartments in the transmission of epigenetic effects (Devanapally et al., 2021; Ewe et al., 2020; Wahba et al., 2021). However, it is still debated whether small RNA-related transgenerational effects transmit via the cytoplasm independently of the nucleus and which heritable agents are involved (see **Discussion**).

Different RNAi factors, which are key for inheritance, for example some of the Argonautes that carry amplified small RNAs, reside in the cytoplasm, while others exist in the nucleus. Argonautes can shuttle between the nucleus and the cytoplasm, and both predominantly nuclear Argonautes, such as HRDE-1 (WAGO-9) and cytoplasmic Argonautes, such as WAGO-4, were shown to be important for RNAi inheritance (Ashe et al., 2012; Buckley B. A. et al., 2012; Luteijn et al., 2012; Shirayama et al., 2012; Xu et al., 2018) (HRDE-1 was recently shown to transfer between the nucleus and cytoplasm, Ding et al., 2023). However, different studies showed that neither Argonaute is absolutely required for RNAi inheritance (Lev et al., 2019b; Ouyang et al., 2022; Xu et al., 2018).

In this work we demonstrate that siRNA-mediated silencing and siRNAs can be inherited in the cytoplasm independently of the nucleus, and investigate the cytoplasmic inheritance mechanism.

## Results

What is the heritable agent that transmits RNAi in the germline to the next generations? Is it RNA (small RNA, mRNA, other), chromatin modifications, or perhaps changes to the chromatin’s architecture? This debate has been ongoing since the discovery of RNAi inheritance, as it is challenging to untangle the different mechanisms from one another. For example, in the nucleus, also in worms, heritable small RNAs guide the trimethylation of histone H3 lysine 9 (H3K9me3), H3K27 and H3K23 (Burton et al., 2011; Gu et al., 2012; Mao et al., 2015; Lev et al., 2017, 2019a), and in multiple organisms it has been shown that chromatin modifications feed-back to induce additional rounds of small RNA synthesis (Grewal, 2023). The capacity of small RNAs to be amplified by RdRPs makes them attractive candidates for the role of the agents which can perpetuate and carry the heritable information transgenerationally without diluting (Rechavi et al., 2011). However, some chromatin marks can similarly self-template (Castel and Martienssen, 2013; Grewal, 2023) (cytosine methylation can famously self-template but the *C. elegans* genome does not contain this modification). The suppressive histone modification that has been most heavily studied in *C. elegans* with regard to RNAi inheritance is H3K9me3, however, the results differed depending on the particular RNAi assay used: While some studies have suggested that H3K9me3 is required for RNAi inheritance (Ashe et al., 2012), later work proposed that the relevant H3K9me3 putative histone methyltransferases (SET-25 and SET-32) are only necessary for the establishment of the silencing response, but not for long term heritable RNAi (Kalinava et al., 2018; Woodhouse et al., 2018). Other studies demonstrated that H3K9me3 is not required at all for heritable silencing of endogenous genes (Kalinava et al., 2017; Lev et al., 2017), and that it might be required only for transgenerational silencing of foreign artificial transgenes, and a small subset of newly evolved genes (Lev et al., 2019a). Further, it is difficult to rule out the possibility that other histone marks are required for dsRNA-induced inheritance in *C. elegans,* and that these other marks compensate for the lack of H3K9me3 in mutants defective in H3K9me3 methyltransferases.

In addition to inheritance of histone modifications, inheritance of other changes to the chromatin, such as the chromatin’s accessibility or condensation (or the chromosomes’ localization in the nucleus), have been demonstrated and thoroughly investigated in different organisms (Wan et al., 2021, 2022). Therefore, also in *C. elegans*, inheritance via changes to the chromatin or other nuclear factors continue to be an important alternative that might explain transgenerational RNAi.

### Detection of nucleus-independent RNAi inheritance using mosaic worms

To test if dsRNA-induced RNAi can be inherited independently of any chromatin changes or the involvement of any other nuclear factor, we utilized an elegant genetic trick that was engineered by Besseling and Bringman (Besseling and Bringmann, 2016). Over-expression of a codon-optimized version of GPR-1, a conserved microtubule force regulator, disrupts the fusion of the maternal and paternal nuclei in the fertilized egg. This leads to uneven segregation of the paternal and maternal nuclei during the first cell division to the blastomeres that will give rise to the germline (P lineage) and the soma (AB lineage). As a result, only one parent contributes its nucleus to the germline of cross progeny of GPR-1-overexpressing (GPR-1(OE)) hermaphrodites. A fluorescently marked strain that was constructed by Artiles et al. (Artiles et al., 2019), enables simple and conclusive identification of such mosaic progeny. Importantly, the entire germline of these chimeric worms derives from a cell with exclusively paternal or exclusively maternal nucleus, and a fused cytoplasm from both origins.

To test for nucleus-independent RNAi inheritance, we exposed GPR-1(OE) mothers to anti-*oma-1* dsRNA, to induce silencing of the redundant germline expressed gene *oma-1* (Detwiler et al., 2001). After selfing, the progeny of these hermaphrodites, which were laid on plates without dsRNA-producing bacteria, were crossed to males carrying a temperature-sensitive and dominant lethal *oma-1(zu405)* allele (Detwiler et al., 2001; Lin, 2003). Silencing of this *oma-1* allele is commonly used to test for RNAi inheritance, because it rescues from embryonic lethality at 20°C (Alcazar et al., 2008; Houri-Ze’evi et al., 2016; Lev et al., 2019b); In other words, only individuals which inherit RNAi survive. In our experiment, we tracked the mosaic progeny which possessed in their germline only the paternally provided nucleus, and therefore their germline genome contained only the *oma-1(zu405)* allele. We found that ∼85% (85.56 ± 6.65 %) of these worms gave rise to viable F3 progeny (**Figure 1**), while empty vector (EV) RNAi treated control worms failed to hatch (3.76 ± 0.18% viable progeny). Since in these experiments the GPR-1(OE) mothers do not transmit their nuclei to the germline of the progeny, we reasoned that the heritable RNAi response was inherited via the ooplasm. Thus, these results reveal that dsRNA-induced RNAi can be inherited in a nucleus-independent manner.

**Figure 1.**
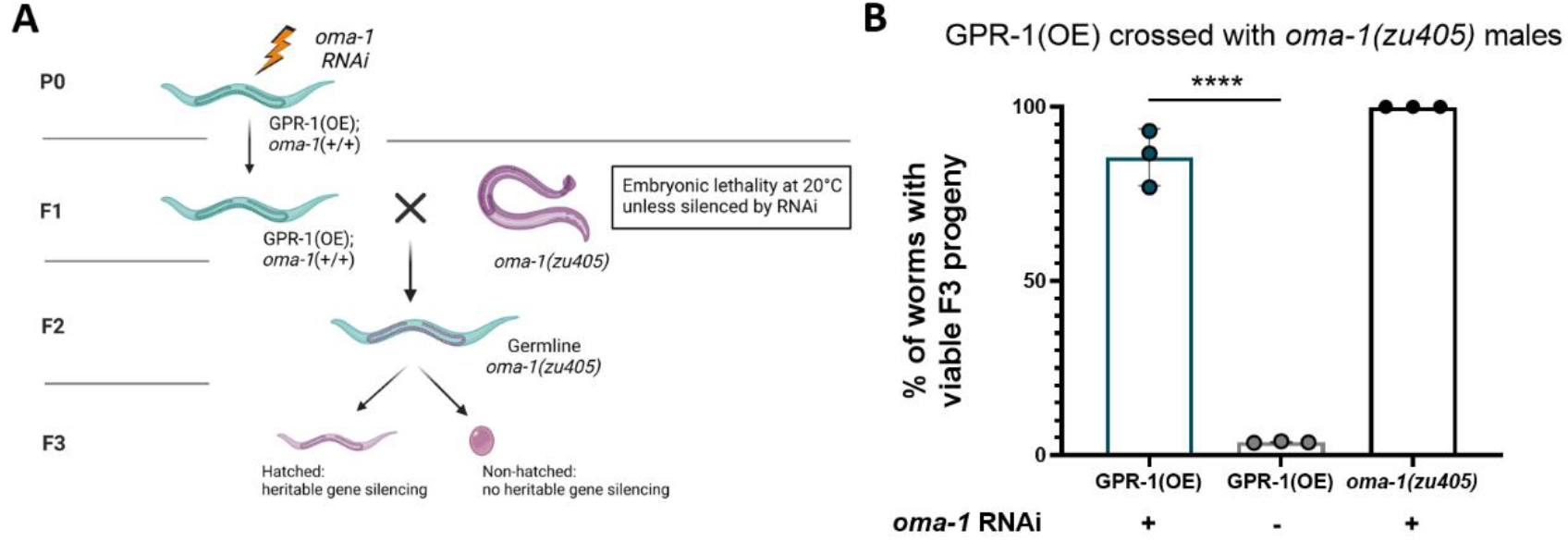
Detection of nucleus-independent RNAi inheritance using chimeric worms. (A) A schematic diagram depicting the experimental procedure. P0 GPR-1(OE) worms were fed bacteria expressing dsRNA complementary to the *oma-1* gene. F1 progeny were then crossed to *oma-1(zu405)* males. ∼80% of chimeric progeny carry paternally derived germline and 2% have a maternally derived germline. We isolated F2 chimeric progeny containing a paternally derived germline and scored for viable F3 progeny. (B) Analysis of nucleus-independent RNAi inheritance. Genotypes on the X axis refer to the P0, dsRNA exposed worms. All worms were crossed to *oma-1(zu405)* males at the F1 generation. *oma-1* RNAi (+) worms were exposed to anti-*oma-1* dsRNA or (-) to control empty vector plasmid. Each dot represents a biological repeat with ∼30 individual F2 worms, bars: mean ± SD. *oma-1(zu405)* (black dots) are the positive control for the RNAi treatment. Additional controls included *oma-1(zu405)* on EV (none of the F1s hatch), F2s with a maternally derived *oma-1*(+) germline (100% of which had viable progeny), data not shown. p value was determined via Fisher’s exact test, **** p-value < 0.0001.

### The nucleus-independent RNAi signal is inherited across generations in the germline and does not result from soma-to-germline transport of dsRNA

In *C. elegans*, RNAi can function non-cell autonomously (Whangbo and Hunter, 2008). Therefore, the silencing response in the paternally derived germline of chimeras could in theory derive from RNA molecules originating in somatic tissues that contain the maternal nuclei. To account for this possibility, we crossed the hermaphrodites to *oma-1(zu405);sid-1(qt9)* mutant males (**Figure 2A**). SID-1 is a transmembrane dsRNA transporter required for import of RNA and systemic RNAi (Winston et al., 2002). In our experiments, the *sid-1(+)* somatic cells are responsive to RNAi and support non-cell autonomous silencing. However, the germline, which contains only the *sid-1(-)* paternal genome, is defective in importing RNAi from cells outside of the germline. We found that SID-1-dependent transport of RNAi from the soma to the germline is not required for nucleus-independent RNAi inheritance (85.44 ± 2.59%), suggesting that the germline already contained sufficient silencing RNA molecules and that the RNAi signal was indeed inherited from the ooplasm (**Figure 2B**).

**Figure 2.**
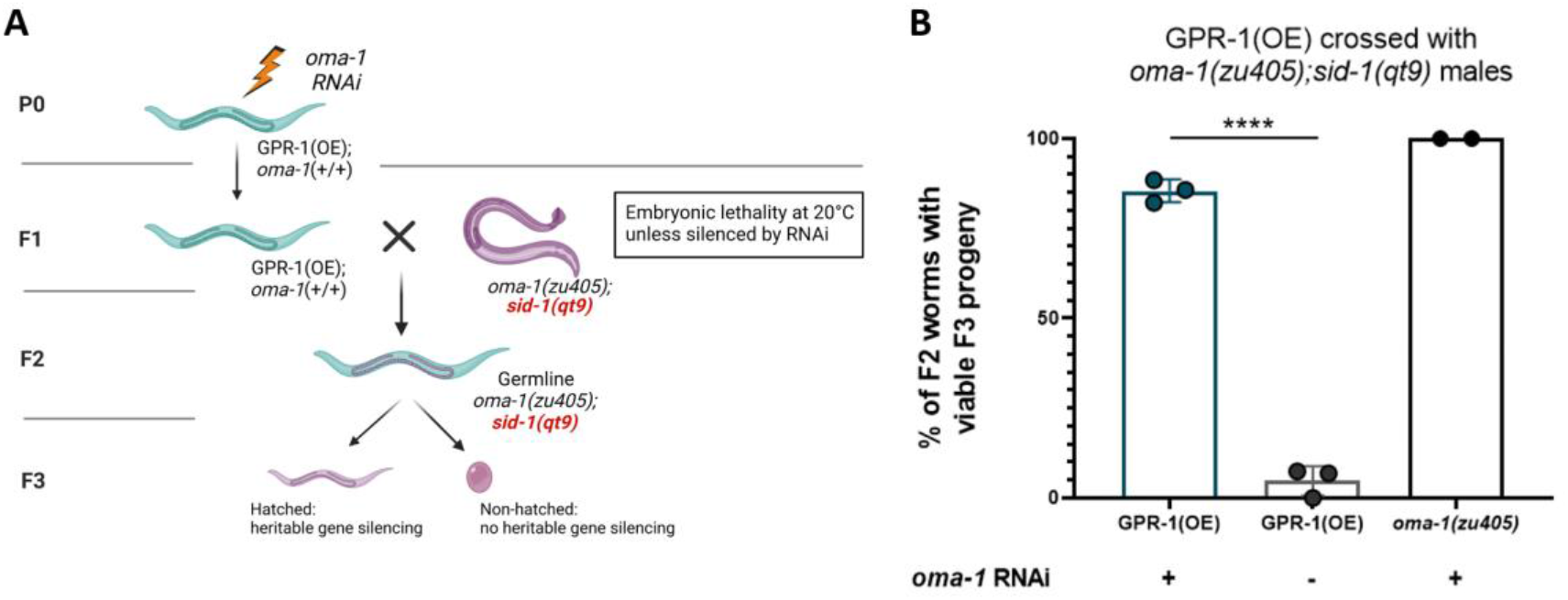
The nucleus-independent RNAi response is inherited across generations in the germline and does not result from soma-to-germline transport of dsRNA. (A) A schematic diagram depicting the experimental procedure. P0 GPR-1(OE) worms were fed bacteria expressing dsRNA complementary to the *oma-1* gene. F1 progeny were then crossed to *oma-1(zu405);sid-1(qt9)* males. F2 chimeric progeny containing paternally derived germline were isolated and scored for viable F3 progeny. (B) Analysis of nucleus-independent RNAi inheritance with *sid-1* mutants. Genotypes on the X axis refer to the P0, dsRNA exposed worms. All worms were crossed to *oma-1(zu405);sid-1(qt9)* males at the F1 generation. *oma-1* RNAi (+) worms were exposed to anti-*oma-1* dsRNA or (-) to control empty vector plasmid. Each dot represents a biological repeat with ∼30 individual F2 worms, bars: mean ± SD. *oma-1(zu405)* (black dots) are the positive control for the RNAi treatment. Additional controls included *oma-1(zu405)* on EV (none of the F1s hatch), F2s with a maternally derived *oma-1*(+) germline (100% of which had viable progeny), data not shown. P-value was determined via Fisher’s exact test, **** p<0.0001.

### ZNFX-1 functions in nucleus-independent RNAi inheritance

Recent evidence highlights the cytoplasmic germ granule component ZNFX-1 as a key player in RNAi inheritance (Ishidate et al., 2018; Wan et al., 2018). It was suggested that ZNFX-1 participates in a small RNA amplification cycle that might function independently of a parallel small RNA amplification cycle that depends on HRDE-1 (Ouyang et al., 2022). We tested whether nucleus-independent RNAi inheritance depends on ZNFX-1. Ouyang et al., found that heritable silencing of *mex-6* and *oma-1* in *znfx-1(gg561)* mutants is weak (partially defective) and therefore concluded that ZNFX-1 and HRDE-1 can compensate for each other (*znfx-1;hrde-1* double mutant were found to be completely defective for RNAi inheritance). Similar redundancy with HRDE-1 was observed for *wago-4* mutants (the Argonaute WAGO-4 colocalizes with ZNFX-1 in the germ granules and the proteins were suggested to work together to promote RNAi inheritance (Wan et al., 2018; Xu et al., 2018)). We successfully replicated the conclusion of these experiments by showing that *znfx-1(gg561)* mutants are capable of partial (weak) RNAi inheritance (24.49 ± 8.91% viable progeny, **Figure 3 third bar from the left**) by testing a different complementary phenotype (see **Methods**). Importantly, we discovered that the observed partial weak RNAi inheritance in *znfx-1* mutants is revoked when inheritance is restricted to the cytoplasm (using the GPR-1(OR) system. 6.87 ± 3.54% viable progeny) (**Figure 3)**. We posited that the residual heritable silencing witnessed in *znfx-1* mutants is mediated by inheritance of epigenetic information in the maternal nucleus and reasoned that ZNFX-1 is involved in nucleus-independent RNAi inheritance.

**Figure 3.**
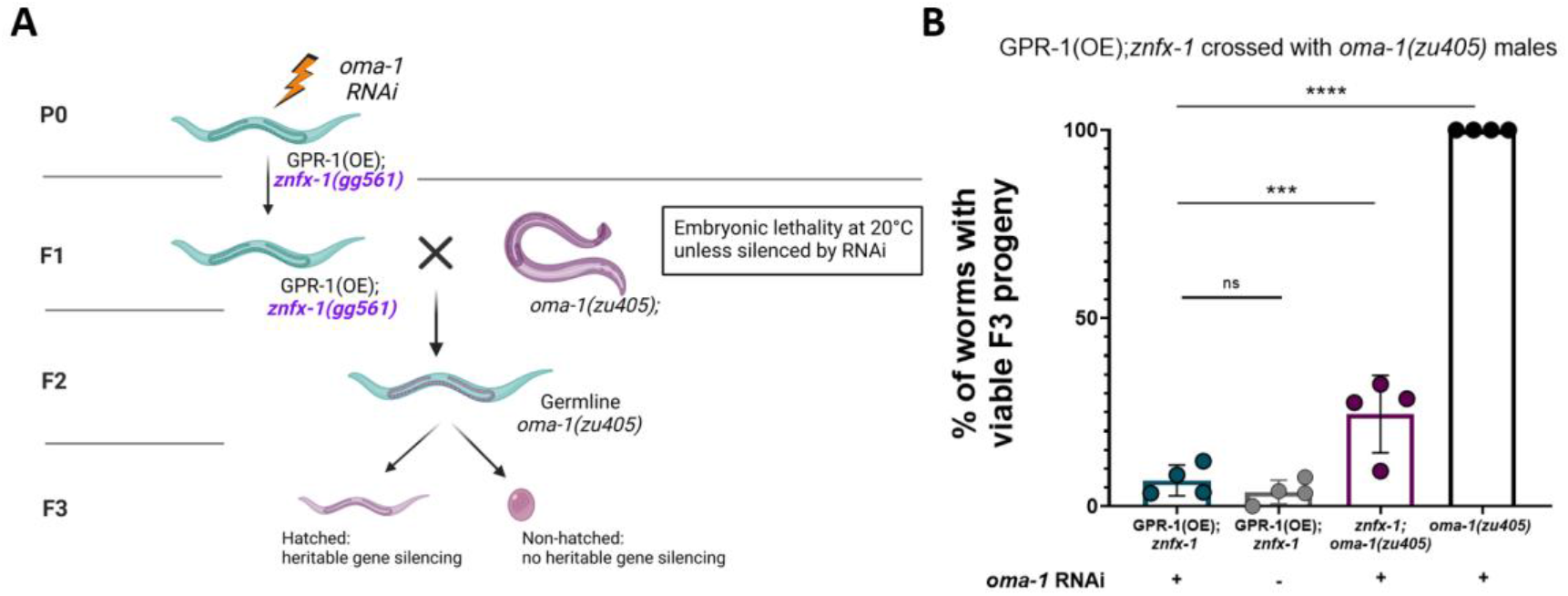
ZNFX-1 is required for nucleus-independent RNAi inheritance. (A) A schematic diagram depicting the experimental procedure. P0 GPR-1(OE);*znfx-1(gg561)* or *oma-1(zu405);znfx-1(gg561)* worms were fed bacteria expressing dsRNA complementary to the *oma-1* gene. F1 progeny were then crossed to *oma-1(zu405)* males. F2 chimeric progeny containing paternally derived germline were isolated and scored for viable F3 progeny. (B) Analysis of nucleus-independent RNAi inheritance in *znfx-1* mutants. Genotypes on the X axis refer to the P0, dsRNA exposed worms. All genotypes were crossed with *oma-1(zu405)* males at the F1 generation. *oma-1* RNAi (+) worms were exposed to anti-*oma-1* dsRNA or (-) to control empty vector plasmid. Each dot represents a biological repeat with ∼30 individual F2 worms, bars: mean ± SD. *znfx-1;oma-1(zu405)* (maroon dots) show partial RNAi inheritance (potentially via the nuclear, *hrde-1* dependent pathway), ***P-value= 0.0003*. oma-1(zu405)*(black dots) are the positive control for the RNAi treatment. Additional controls included *oma-1(zu405)* on EV (none of the F1s hatch), F2s with a maternally derived *oma-1*(+) germline (100% of which had viable progeny), data not shown. P-values were determined via Fisher’s exact test, **** P-value<0.0001 – indicates no inheritance in *znfx-1* mutation background to GPR-1(OE).

### Germ granules disruption in *pptr-1* mutants potentiates nucleus-independent RNAi inheritance and enables transgenerational silencing even in *znfx-1* mutants

Which cytoplasmic components or organelles are involved in nucleus-independent RNAi inheritance? Since ZNFX-1 localizes to cytoplasmic germ granules, we reasoned that germ granules function in nucleus-independent dsRNA-mediated siRNA inheritance. The germ granules are deposited from the ooplasm to the P lineage in the early embryo (Strome and Wood, 1983). While the granules are important for RNAi and for RNAi inheritance (Lev and Rechavi, 2020; Sundby et al., 2021), it was recently shown that RNAi can nevertheless be inherited even when the deposition of germ granules to the embryo is severely disrupted (Dodson and Kennedy, 2019; Lev et al., 2019b). To examine the role of proper germ-granule segregation to cytoplasmic inheritance, we tested nucleus-independent RNAi inheritance in *pptr-1* mutants (BFF131: GPR-1(OE);*pptr-1(tm3103)*). PPTR-1 encodes for a regulatory subunit of the conserved phosphatase PP2A, and is necessary for the correct, asymmetrical segregation of P granules to the embryo’s P lineage and ultimately to the germline. In *pptr-1(tm3103)* mutants, P granules are incorrectly segregated and disassembled at each division, resulting in germ cells containing P granules that are reduced in both size and number (Gallo et al., 2010).

Previously we found that, surprisingly, heritable RNAi silencing is much stronger in *pptr-1(tm3103)* mutants compared to wild-type worms (Lev et al., 2019b). The effect is dramatic, as heritable RNAi responses last 3-5 generations on average in wild type worms (Alcazar et al., 2008; Houri-Ze’evi et al., 2016), while in *pptr-1* mutants they last for at least 70 generations (Lev et al., 2019b). Here we show that when *pptr-1* is disabled, nuclear-independent RNAi inheritance is potentiated and lasts longer (**Figure 4A-C**). Further, we found that when nuclear-independent RNAi inheritance is potentiated in *pptr-1* mutants, RNAi is weakly inherited via the cytoplasm even in the absence of ZNFX-1 (19.25 ± 10.94% viable progeny) (**Figure 4D**).

**Figure 4.**
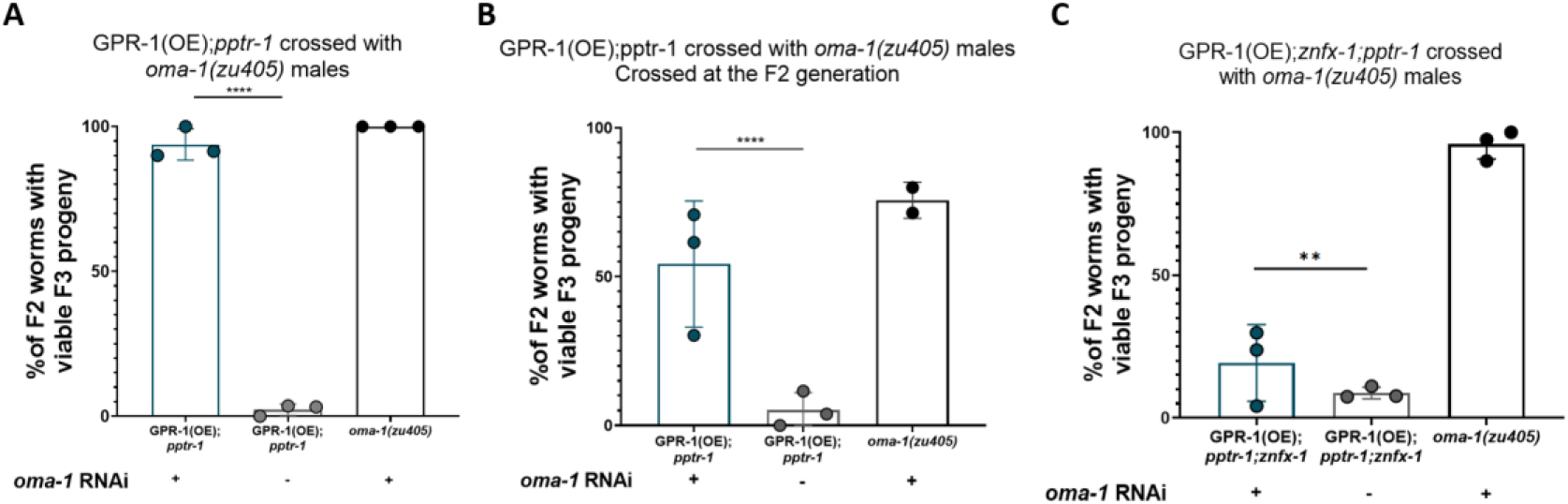
The involvement of germ granules in nucleus-independent RNAi inheritance. Analysis of nucleus-independent RNAi inheritance in *pptr-1*mutants (A,B) or *pptr-1;znfx-1* (C). Genotypes on the X axis refer to the P0, dsRNA exposed worms. All genotypes were crossed with *oma-1(zu405)* males at the F1 generation (A,C) or F2 generation (B). *oma-1* RNAi (+) worms were exposed to anti-*oma-1* dsRNA or (-) to control empty vector plasmid. Each dot represents a biological repeat with ∼30 individual F2 (A,C) or F3 (B) worms, bars: mean ± SD. *oma-1(zu405)* (black dots) are the positive control for the RNAi treatment. Additional controls included *oma-1(zu405)* on EV (none of the F1s hatch), F2s (A,C) or F3s (B) with a maternally derived *oma-1*(+) germline (100% of which had viable progeny), data not shown. P-values were determined via Fisher’s exact test, **** p<0.0001 – indicates inheritance in *pptr-1* mutation background to GPR-1(OE), ** p=0.0094– indicates inheritance in *pptr-1;znfx-1* mutation background to GPR-1(OE).

Altogether, these results suggest that the fidelity of germ granule segregation determines the strength of cytoplasmic siRNA inheritance in a ZNFX-1-dependent manner, however, ZNFX-1 is not absolutely required for cytoplasmic inheritance.

### Endogenous siRNAs are inherited in a nucleus-independent manner

Next, we tested if endogenous small RNAs are also inherited via the cytoplasm, similarly to exogenous dsRNA-derived siRNAs. To do so we crossed different *C. elegans* isolates. We crossed hermaphrodites of the lab strain N2 (isolated in Bristol, UK) which over-express GPR-1, with MY16 males (isolated in Munster, Germany). These two isolates differ in 111,533 single nucleotide polymorphisms (SNPs) and 17,410 indels across the genome (Cook et al., 2017) and moreover differentially express ∼1,850 endogenous small RNAs (**Sup Figure 1**). By combining the non-mendelian inheritance of GPR-1(OE) mutants and the polymorphism between the two isolates we were able to search for endogenously derived heritable endogenous small RNAs: Over expression of GPR-1 in N2 resulted in chimeric F1 worms that gave F2 offspring that were genetically identical to either MY16 or N2 (**Figure 5A**), which we refer to as F2 MY16* and F2 N2*, respectively. After crossing the worms, we sequenced small RNAs and mRNAs from multiple biological and technical replicates of F2 N2, F2 N2*, F2 MY16 and F2 MY16* worms. Sequencing and analysis of mRNAs confirmed that indeed the nuclear mRNAs from F2 MY16* worms corresponded to the MY16 genome, while the cytoplasmic, mitochondrially encoded mRNAs in MY16* corresponded to the N2 mitochondrial DNA.

**Figure 5.**
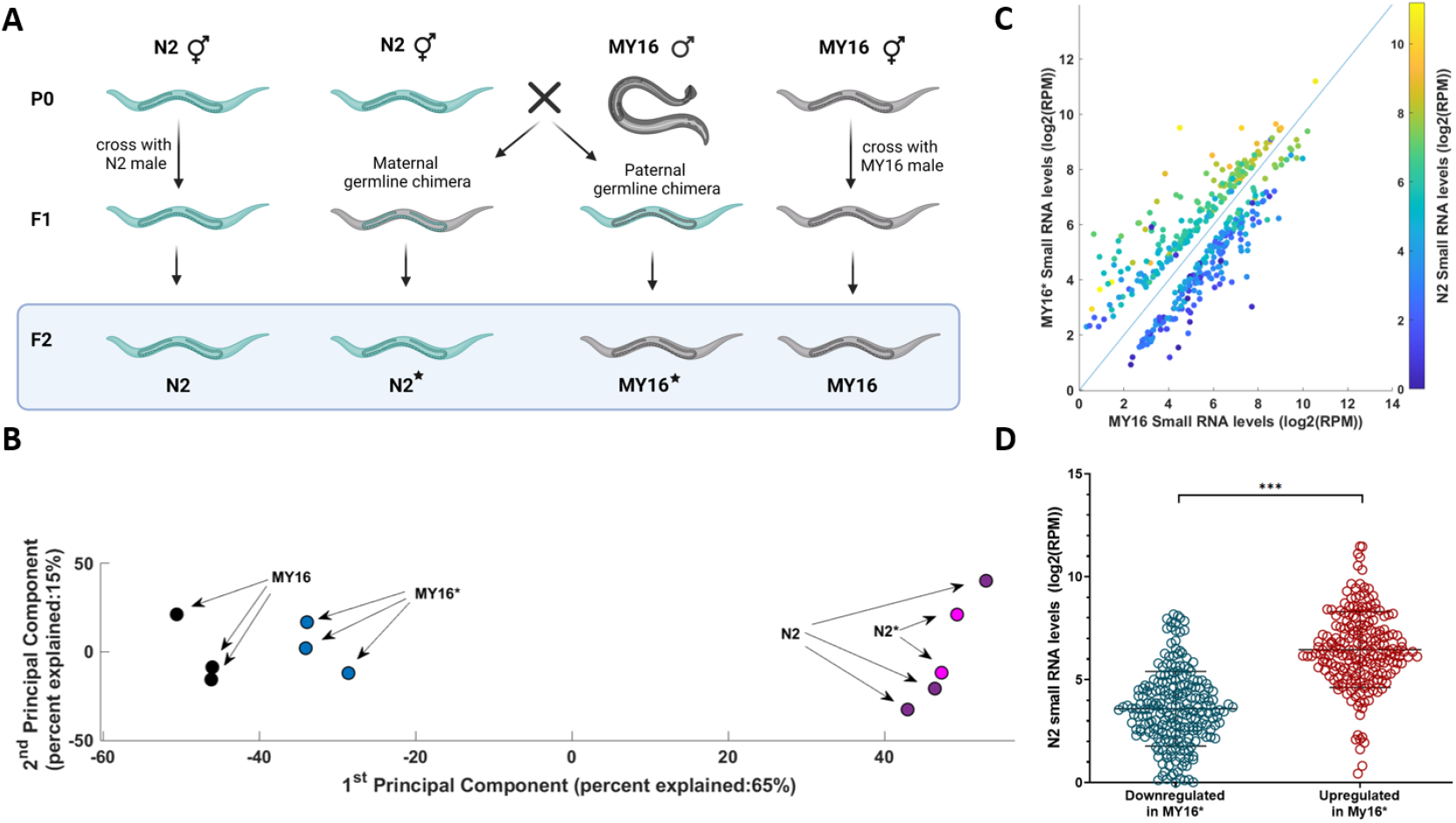
Endogenous siRNAs are inherited in a nucleus-independent manner. (A) A schematic diagram depicting the experimental procedure. P0 N2 worms, over-expressing GPR-1, were crossed to MY16 males (middle) to generate mosaic F1 progeny with either maternal or paternal germlines. F2 progeny coming from F1 with paternal germlines are genotypically MY16, and are named MY16*, while F2 progeny of F1 worms with maternal germlines are genotypically N2 and are named N2*. In addition, P0 N2 worms, over-expressing GPR-1, were crossed to N2 males to ultimately generate F2 N2 (left) and MY16 worms were crossed to MY16 males to generate F2 MY16 worms. Small RNA and mRNA libraries were generated from the F2 generation. (B) A PCA projection of small RNA targeting protein coding genes from 4 conditions. MY16*and MY16 are located distinctly, with MY16* closer to N2 and N2* that MY16 is. The % variance, out of the total original variance in the high-dimensional space, spanned by the first and second PCs is indicated on the x and y axis, respectively. (C) Comparison of small RNA expression levels (log2 of reads per million (RPM)) of genes that are differentially expressed between MY16 worms (X axis) and MY16* worms (Y axis), with color code indicating the corresponding small RNA levels in N2 worms. (D) Comparison of genes that are downregulated in small RNA reads in MY16* worms relative to MY16 (left) and upregulated (right). Y axis indicates small RNA levels in N2 worms. Genes that are upregulated in MY16* worms have significantly higher expression of small RNA levels in N2 worms than genes that are downregulated, suggesting inheritance from P0 N2 to F2 MY16* (Wilcoxon rank sum test, p-value = 5.1775e-40).

We detected cytoplasmic inheritance of endogenous small RNAs that target 129 protein-coding genes (these small RNAs are transmitted via the cytoplasm of P0 N2 to F2 MY16*). We determined that these endogenous small RNAs were indeed inherited via the mother’s cytoplasm based on three observations: (1) The small RNA pool of MY16* worms exhibited higher similarity to the small RNA pool of N2 worms than did the small RNA pool of MY16, based on principle component analysis (PCA) (**Figure 5B**). (2) We expected that accumulation of heritable small RNAs would increase the levels of these RNAs in the worms that inherit them, and indeed, the small RNAs that were up-regulated in MY16* compared with MY16, had higher levels in N2 worms than the small RNAs that were down-regulated in MY16* worms compared with MY16 worms (**Figure 5C, D**). (3) We found that small RNAs that were up-regulated in MY16* were enriched for ZNFX-1 class small-RNAs (see **Methods**) (fold enrichment of 6.2, P<0.0001) consistent with the results we describe above regarding inheritance of exogenous RNAi responses. Further, we found that the cytoplasmically inherited endo-siRNAs that transmit from the N2 isolate to the MY16 progeny are enriched for siRNAs that target genes that are poorly conserved between *Caenorhabditis* species (fold enrichment of 1.95, P=0.0019).

Which types of RNA molecules are inherited in the germ granules? Unlike 22G siRNAs which can be amplified by RdRPs, mRNAs should get diluted and are therefore not expected to be inherited in large enough quantities to enable information transfer beyond the F1 generation. Still, one might imagine a scenario in which hypothetical “heritable mRNAs” are concentrated in germ granules so that enough parentally transcribed mRNA molecules get deposited in the oocytes and remain functional even after more than one generation (namely, enough remain to serve as templates for RdRP-mediated de novo synthesis of 22G amplified siRNAs that would lead to silencing). To examine this possibility, we analyzed SNPs that differentiate N2 and MY16 animals. Mitochondrial RNA in MY16* contained SNPs that match the N2 genome, as MY16* worms inherited their mitochondria from their N2 grandmothers. In contrast, we found that nuclearly encoded mRNAs in MY16* match the MY16 genome, and not N2 genome; In other words, we could not detect statistically significant levels of N2 nuclear SNPs in RNAs that were inherited by MY16* worms. We reasoned therefore that the majority of the heritable cytoplasmic RNAs are not mRNAs but instead 22G siRNAs which are amplified again and again in every generation based on newly transcribed nuclear mRNA templates (and not based on miniscule amounts of mRNAs that avoid dilution and get inherited via the cytoplasm). This suggests that the heritable agents, the RNA molecules which are inherited in the in the germline, are siRNAs, which are capable of RdRP-mediated amplification, and not granules-enriched mRNA templates.

## Discussion

Perturbing the small RNA pool or the germline chromatin landscape leaves a transgenerational trace in the progeny. Our results suggest that cytoplasmic inheritance of small RNA-mediated silencing can occur independently of chromatin modifications or other nuclear factors for multiple generations.

Previous studies examined the dependency of different heritable epigenetic effects on the nucleus, and reached different conclusions depending on the assay and phenotype tested. For example, starvation-induced effects on gut development were found to be transmitted by nuclear factors, independently of heritable cytoplasmic maternal factors (Ewe et al., 2020). Interestingly, many mutants defective in both cytoplasmic and nuclear factors (both in small RNA and in chromatin genes) display a transgenerational loss of fertility phenotype termed “Mortal Germline” (Mrt) (Ahmed and Hodgkin, 2000; Batista et al., 2008; Katz et al., 2009; Zhang et al., 2011; Conine et al., 2013; Simon et al., 2014; Xu et al., 2018; Manage et al., 2020; Schreier et al., 2022). Recently, Wahba et al. suggested that sterility factors which lead to a Mrt phenotype pass exclusively via the cytoplasm (Wahba et al., 2021). Wahbe et al. proposed that these cytoplasmically inherited sterility factors could be small RNAs that regulate ribosomal RNAs. Another study argued that other silencing factors could also have important contributions for the Mrt phenotype (Montgomery et al., 2021) (it was not directly examined where in the cell do these factors localize). A different heritable phenotype, termed mating-induced transgene silencing, was also shown to be nucleus-independent (Devanapally et al., 2021). Since our results show that siRNAs can transmit cytoplasmically without the nucleus, it strengthens the possibility that different heritable epigenetic phenotypes, such as Mrt and mating-induced silencing, result from aberrant cytoplasmic inheritance of siRNA molecules.

The germ granules organize the production of heritable small RNAs and are crucial for cytoplasmic RNAi inheritance. Our results show that *znfx-1* mutants are defective in RNAi inheritance via the cytoplasm; However, disrupting proper segregations of the granules in *pptr-1* mutants paradoxically strengthens dsRNA-mediated cytoplasmic RNAi inheritance, and enables weak cytoplasmic RNAi inheritance even in *znfx-1* mutants. One possible explanation could be that this potentiation of cytoplasmic inheritance occurs since in *pptr-1* mutants other inherited endogenous small RNAs are not properly synthesized, rendering more small RNA-biogenesis machinery free to produce exogenous siRNAs (as hypothesized in (Lev et al., 2019b)). Competition between different small RNA species has been raised as an explanation for transgenerational effects multiple times in the past (Houri-Zeevi et al., 2020; Houri-Ze’evi et al., 2016; Sarkies et al., 2013; Zhuang and Hunter, 2012), and an increase in the levels of exogenous small RNAs, coupled with a decrease in the levels of endogenous small RNAs has been indeed documented in *pptr-1* mutants (Lev et al., 2019b).

A recent publication by Schreier et al. (Schreier et al., 2022) described epigenetic inheritance of the Mrt phenotype via *C. elegans* males. This inheritance was found to be mediated by the Argonaute protein WAGO-3, which is associated with 22G RNAs and is expressed in sperm cells. Additionally, Schreier et al. identified and defined the presence of Paternal Epigenetic inheritance (PEI) granules, which are specific to sperm cells and get inherited along with WAGO-3. Our sequencing data did not reveal inheritance of cytoplasmic RNAs from sperm (see **Figures 6A,B, Sup Figure 2**), perhaps because the cytoplasmic content of the sperm is very small relatively to the oocyte’s. It would be interesting in the future to study the pool of paternally inherited endogenous small RNAs.

Small RNA inheritance in *C. elegans* requires amplification of 22G siRNAs using RdRPs. However, small RNAs are not amplified and inherited forever, as it appears that small RNAs cannot themselves serve as templates for amplified small RNA synthesis (Pak et al., 2012). Our analysis of RNA sequencing and SNP data strengthens the hypothesis that heritable small RNAs require de novo synthesis of mRNA templates as neither significant amounts of P0 parental small RNAs nor mRNA templates were found to persist all the way to the F2 generation. This could be a fundamental limit on the independence of RNA encoded information from DNA.

“Epigenetic reprogramming” is one of the major theoretical boundaries preventing epigenetic transmission of parental responses across generations (Heard and Martienssen, 2012). This process entails the resetting of almost all epigenetic information in the nucleus, including both DNA and histone modifications, presumably so that the next generation can start as a “blank slate”. Importantly, work in some organisms, notably fish, shows that chromatin reprogramming is not absolutely required, (Skvortsova et al., 2019). Studying RNAi-triggered epigenetic inheritance in *C. elegans* is technically easy, and thus continue to provide new insights into the mechanisms of heredity, including the revelation that gene regulatory responses can transmit via cytoplasmic small RNAs transgenerationally even when the entire nucleus is replaced, regardless of whether the chromatin epigenetic information is modified or “reprogrammed”.

## Acknowledgments

We thank all the Rechavi and Shaham lab members for their helpful comments and fruitful discussions. We thank Prakash Cherian and Omri Wurtzel for their assistance. We are grateful to Itai A. Toker for his comments and suggestions. Some strains were provided by the CGC, which is funded by NIH Office of Research Infrastructure Programs (P40 OD010440). Some of the figures in this article were created with BioRender.com. This work was supported by ERC grant #33562. I.R is supported partly by a fellowship from the Prajs-Drimmer Institute. O.R is grateful for funding from the Eric and Wendy Schmidt Fund for Strategic Innovation (Polymath Award #0140001000) and the generous funding from the Khan foundation (Grant # 0604918421).

## Methods

### Cultivation of worms

All the experiments were performed at 20 degrees Celsius, except for maintenance of strains containing the *oma-1(zu405)* allele, which was done at 15 degrees Celsius. Worms were cultivated on NGM plates seeded with OP50 bacteria apart from when treated with HT115 bacteria that express dsRNAs for RNAi induction.

These strains were used in this work: PD2218 ccTi1594 [mex-5p::GFP::gpr-1::smu-1 3’UTR + Cbr-unc-119(+), III: 680195] III. umnIs7 [myo-2p::GFP + NeoR, III:9421936] III, YY996 znfx-1(gg561), TX20 oma-1(zu405), JH2787 pptr-1(tm3103), HC196 sid-1(qt9), MY16, N2 and several combinations made by us by crossing strains and validating genotype using PCR and/or Sanger sequencing.

The nematodes were kept well fed and extra care to avoid contamination was taken. Contaminated or starved plates were discarded and not analyzed.

### GPR-1(OE) crosses

Hermaphrodites carrying both ccTi1594 and umnIs7 transgenes, containing GPR-1(OE) and a pharyngeal marker, respectively, were crossed to *oma-1(zu405)* or *oma-1(zu405);sid-1(qt9)* males. Non-mendelian progeny of these crosses were identified using the pharyngeal fluorescent marker, to identify F2 chimeric worms containing maternal chromosomes in the AB cell lineage and paternal chromosomes in the P1 cell lineage as described by Artiles et al. (Artiles et al., 2019). As C. elegans’ germline derives entirely from the P1 cell lineage (Sulston et al., 1983), the worms’ germline contained the temperature sensitive *oma-1(zu405)* allele. OMA-1 silencing was quantified by scoring the number of worms that lay five or more viable progeny, as previously described (Houri-Ze’evi et al., 2016).

### RNAi Treatment

HT115 bacteria that transcribe dsRNA targeting *oma-1* or control empty-vector that does not lead to dsRNA transcription and gene silencing were grown in LB with 100 μg/ml carbenicillin. Bacteria were then seeded on NGM plates that contain Carbenicillin (25 μg/ml) and IPTG (1 mM). Worms were put on RNAi plates 24 hours after seeding for two generations. First generation was put at the L4 stage. Worms were treated on RNAi plates for 2 consecutive generations. Before transferring worms from RNAi plates to NGM plates, the worms were washed 4 times in M9 buffer to remove dsRNA-inducing bacteria.

### Worm collection and RNA extraction

Hermaphrodites were collected on the first day of adulthood, washed 4 times in M9 buffer and collected into an Eppendorf tube prior to the addition of 4 volumes of Trizol (Life Technologies) to 1 volume of worm/M9 pellet. To extract RNA the following protocol was used: 3 freeze/thaw cycles - −80 °C for 30 minutes followed by 15 minutes vortex at room temperature. 1 volume of chloroform was added to 5 volumes of Trizol/worm/M9. The mix was put in a pre-spun 2mL Heavy Phase-Lock tube and centrifuged at 16000 g for 5 minutes at 4 °C. The aqueous phase was transferred to new pre-spun 2mL Heavy Phase-Lock tube, and 1 volume of Phenol:Chloroform:Isoamyl alcohol was added per 1 volume of aqueous phase. The tube was centrifuged at 16000 g at room temperature. Aqueous phase was transferred to Eppendorf tube and precipitated with 1 volume of isopropanol and 1.3 µL glycogen (20 µg/µL). the tubes were put for 30 minutes at −20 °C before centrifugation at 16000 g at 4°C. Pellet was washed with 900µL of cold 70% EtOH, and left at room temperature for 20 minutes before being left at −20°C over night. The next morning, the tube was centrifuged at 16000 g for 10 minutes, and the pellet was washed again with cold 70% EtOH. Tubes were centrifuged at 16000 g for 10 minutes, all EtOH was removed and pellet was resuspended in 12 µL of warm (70°C) ddH2O. RNA concentration was determined using Qbit and RNA quality was tested using Agilent 2200 TapeStation.

To ensure the capture of small RNA regardless of their 5’ phosphorylation status, 150-1000 ng from each sample was treated with RNA 5’ polyphosphatase. Concentrations and quality were assessed using Qubit and Agilent 2200 TapeStation respectively.

### sRNA libraries

The NEBNext® Multiplex Small RNA Library Prep Set for Illumina® from New England Biolabs® was used for small RNA library preparation, following the manufacturer’s protocol. The concentration of the samples was determined using Qubit, and their quality was assessed using an Agilent 2200 TapeStation. Subsequently, the samples were pooled and electrophoresed on a 4% agarose E-Gel from Life Technologies. Bands ranging in length from 140 to 160 nt were carefully excised and purified using the MiniElute Gel Extraction Kit (QIAGEN). The purified samples were once again assessed for quality and concentration using the Agilent 2200 TapeStation. Libraries were sequenced on an Illumina NextSeq500 sequencer.

### mRNA libraries

The NEBNext® Ultra II Directional RNA Library Prep Kit for Illumina® from New England Biolabs® was used for mRNA library preparation, following the manufacture’s protocol. The concentration of the samples was determined using Qubit, and their quality was assessed using an Agilent 2200 TapeStation. Samples were then pooled together prior to sequencing on an Illumina NextSeq500 sequencer.

### Small RNA seq analysis

The Illumina *.fastq output files were first assessed for quality, using FastQC (Simon Andrews, 2010). The files were then assigned to adapters clipping using Cutadapt (Martin, 2011). Next, the clipped reads were aligned against the ce11 version of the C. elegans genome using ShortStack (Shahid and Axtell, 2014). We counted reads which align in antisense orientation to genes, using the python-based script HTSeq-count (Anders et al., 2014) and the Ensembl-provided gff file (release-95).

We then assigned the summarized counts for differential expression analysis using the R package DESeq2 (Love et al., 2014) and limited the hits for genes which were shown to have an FDR < 0.1.

### ZNFX-1-class small RNAs

to generate a list of ZNFX-1 regulated genes, we analyzed available data from Ouyang et al. (Ouyang et al., 2022) as written above. ZNFX-1 regulated genes are genes targeted by small RNAs that were downregulated in *znfx-1* mutants compared to wild-type N2.

### mRNA seq analysis

mRNA libraries were first assessed for quality using the FastQC tool (Andrews, 2010) and were then aligned to ce11 version of the genome using HISAT2 (Kim et al., 2015). The aligned reads were then counted using the python-based script HTSeq-count (Anders et al., 2014) and the Ensembl-provided gff file (release-95). Next, the samples were then compared for differential expression using the R package DESeq2 (Love et al., 2014). Genes were regarded as differentially expressed if they pass the criterion of FDR < 0.1.

### SNP analysis

We first aligned the FASTQ files from small RNA sequencing of N2 and MY16 isolates to the ce11 reference sequence, while allowing up to two mismatches. We pooled the aligned reads of each strain, and calculated the frequency of each of the four nucleotides in each covered position. We only included reads with length > 20 nucleotides, which are uniquely aligned to the genome.

For a given genomic position, we defined the occurrence of SNP if (1) it was covered by at least 5 reads per million in each isolate, (2) it was dominated (frequency > 0.95) by specific nucleotide in each isolate, and (3) the identity of the dominant nucleotide differs between N2 and MY16. We further compared the SNPs defined by us to the SNPs reported in CeNDR, and found that more than 80% of the SNPs detected by us also appear in the CeNDR database.

Next, we aligned the FASTQ files from small RNA sequencing of MY16* to the ce11 reference genome under the same conditions, and looked for cases in which the identity of the nucleotide in each given position, in at least 5% of the occurrences, is deviates from that of the MY16 isolate, but identical to that of the N2 strain. In order to avoid the possible effect of sequencing error and other biases, we further demand that the representations of the N2-like nucleotide among the nucleotides covering a given position in the MY16* isolate will be higher by at least 1.5-fold from the representations of the N2-like nucleotide observed for the MY16 one. Finally, in order to evaluate the significance of our results, we ran a similar analysis, in which the definition of SNPs was determined using the N2 and MY16* strains, while looking for cases in which the identity of the MY16’s nucleotide at a given position follows that of N2 and deviates from the MY16*.

## Statistical Analyses

Statistical analyses and generation of graphs were performed using Prism, R v4.0.0. Statistical details of experiments appear in figure legends.

For viable progeny analysis, we used Fisher’s exact test, with two-stage step-up method of Benjamini, Krieger, and Yekutiely correction for multiple comparison.

## Decleration of interests

The authors declare no competing interests

## Supplementary Figures

**Supplementary Figure 1.**
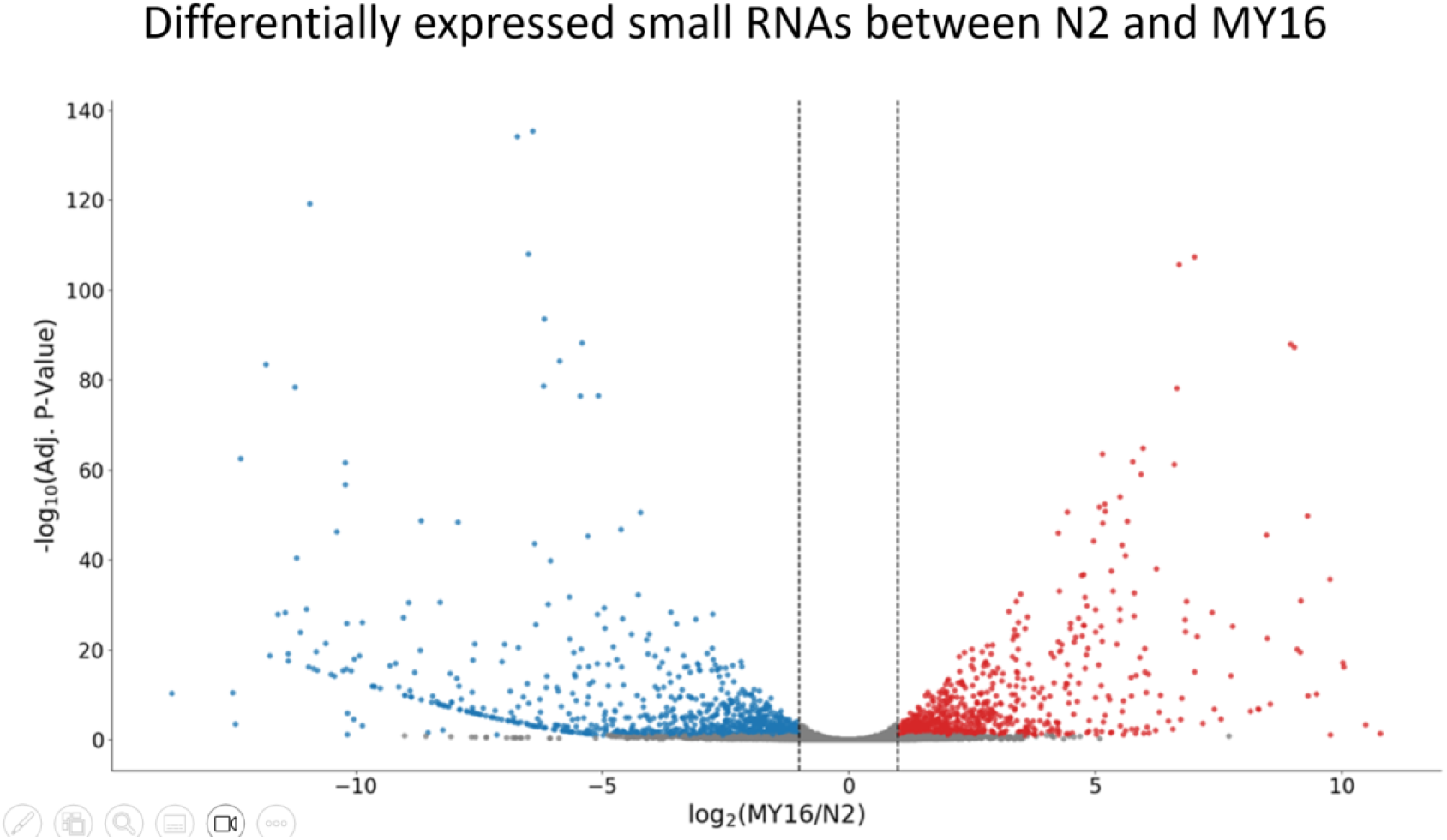
Differentially expressed small RNAs between N2 and MY16. (A) volcano plot for small RNA data for all genes. Genes that are significantly upregulated in MY16 compared to N2 are colored red while genes that are significantly downregulated in MY16 compared to N2 are colored red. Y axis represents statistical significance.

**Supplementary Figure 2.**
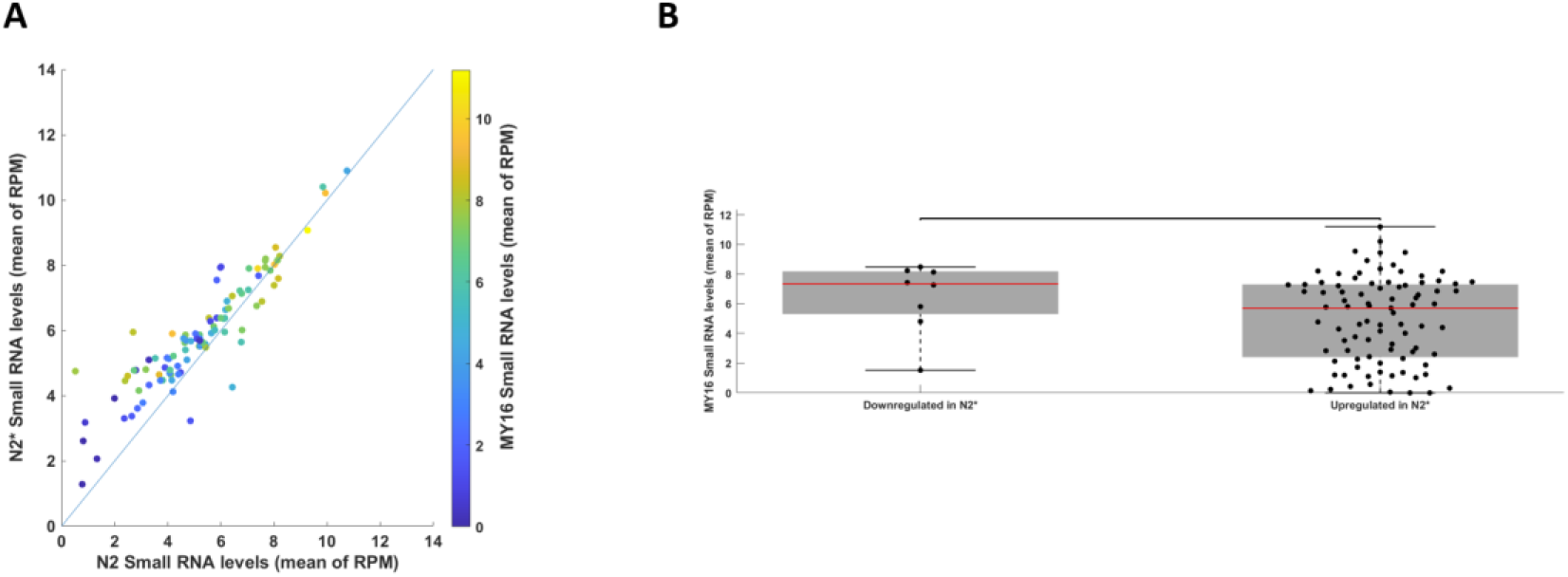
no detection of small RNA inheritance from sperm’s cytoplasm. (A) Comparison of small RNA expression levels (log2 of reads per million (RPM)) of genes that are differentially expressed between N2 worms (X axis) and N2* worms (Y axis), with color code indicating the corresponding small RNA levels in MY16 worms (B) Comparison of genes that are downregulated in small RNA reads in N2* worms relative to N2 (left) and upregulated (right). Y axis indicates small RNA levels in MY16 worms. There is no significant difference between the two sets of genes.

